# Autophagy is associated with survival and resiliency of fibroblasts in long-term decaying post-mortem parenchymal tissue

**DOI:** 10.1101/2022.08.18.504466

**Authors:** Kathleen M. McAndrews, Daniel J. McGrail, Liv B. Gansmo, Sara P.Y. Che, Deepavali Chakravarti, Valerie S. LeBleu, Raghu Kalluri

**Author notes:** Correspondence: Raghu Kalluri, MD, PhD,.

## Abstract

Fibroblasts are spindle-shaped mesenchymal cells and an abundantly studied cell type that are easy to culture. Their adaptive response in culture conditions allows for use in many different cell biological experiments, including their utility in generating induced pluripotent stem cells. Despite extensive use of fibroblasts in cell and molecular biology and genetics experiments, fundamental evaluation of their resiliency and survival programs, in comparison with other cell types, is undetermined. Here, we demonstrate that fibroblasts exhibit remarkable survival capacity in post-mortem tissue decaying at room temperature and can be cultured from ear, tail, kidney, lung, fetal, and mammary tumor tissue after 12-hours of post-mortem tissue decay. Fibroblasts can be cultured from ear and lung tissue after 24-hours, and from ear after up to 120-hours of post-mortem tissue decay. Gene expression profiling of post-mortem lung tissue fibroblasts compared to fresh tissue cultured fibroblasts suggested a transition to a more quiescent phenotype with activation of nutrient scavenging pathways as evidenced by downregulation of genes associated with DNA replication, ribosomes, cell cycle, and spliceosomes as well as upregulation of genes associated metabolism, autophagy, and lysosomes. Measurement of light chain 3B (LC3B)-I/LC3B-II ratio and lysosomal-associated membrane protein (LAMP)-1 indicate that autophagy is increased in post-mortem fibroblasts, with evidence for potential increase in autolysosomes and senescence program. Our study provides evidence for the ability of normal fibroblasts to overcome extreme stress conditions and offers new insights into cell survival mechanisms and aging, with potential utility in tissue regeneration and repair.

## Introduction

Fibroblasts are mesenchymal cells with critical roles in embryonic development, and the formation and maturation of the connective tissue [1, 2]. Fibroblasts likely also play an important role in parenchymal organogenesis [3-5]. In adult parenchymal organs, fibroblasts regress to minimal numbers but can expand in the context of tissue repair and regeneration [6], organ fibrosis [7], and cancer [8]. Fibroblasts produce extracellular matrix molecules including the most abundant human protein, type I collagen [9, 10]. Type I collagen produced by fibroblasts is speculated to play a role in tissue fibrosis [11], cancer progression and metastasis [12].

Fibroblasts were one of the first cell types to be cultured on a plastic dish due to their ease of cultivation [13-15]. Cultured fibroblasts from embryonic tissue and skin are commonly used in numerous cell and molecular biology experiments to discover new signaling pathways and identify new proteins [16]. A prototype of embryonic fibroblasts is the NIH 3T3 cells, isolated by George Todaro and Howard Green from a Swiss Albino mouse [15]. The NIH 3T3 cell line was extensively used in many cell culture experiments due to their ability to grow on a plastic substrate and their high proliferative index. It was subsequently shown that NIH 3T3 were not ‘normal’ fibroblasts, as they exhibit chromosomal aberrations [15, 17, 18], and in some contexts, could form tumors [19]. Many established fibroblast cell lines commonly used in laboratory research are genetically abnormal [18, 20, 21] and are therefore more likely to be efficiently sustained in culture. In this regard, fibroblasts from disease tissue expand more readily grow in culture than fibroblasts from normal counterpart tissue, and epigenetics is speculated to play a role in this differential culture adaptive response [22-25].

Fibroblasts isolated from mouse and human dermis or human neonatal foreskin are commonly used as precursors to generate induced pluripotent stem cells (iPSCs) [26]. Normal fibroblasts are considered to have low epigenetic drifts [27] and can withstand manipulation associated with transfection reagents and viral infection to generate iPSCs [26]. In addition, they are known to express low levels of pluripotency genes such as *Oct3/4, Sox2, Nanog* and *Klf4* [28]. Anecdotal reports have been published which suggest that post-mortem human scalp derived fibroblasts from frozen cadavers [29] and post-mortem human skin fibroblasts from a rapid autopsy (3-7 hours post-mortem) [30] can be used to generate iPSCs.

Despite extensive research on fibroblasts, their in vivo resiliency under stress conditions remains unknown. In this report we used severe stress (post-mortem decay at room temperature) to ascertain the survival and resiliency programs of fibroblasts from multiple tissues and identify putative mechanism associated with such unique features of fibroblasts.

## Results

### Live fibroblasts can be isolated from post-mortem tissue

To determine whether fibroblasts can be isolated from mouse tissue of distinct origins, we utilized alpha smooth muscle actin (αSMA) promoter-driven red fluorescence protein (αSMA-RFP) mice. These mice allow for flow cytometry analysis and identification of a class of fibroblasts referred to as myofibroblasts which are characterized by αSMA expression in culture [8, 31, 32]. Employing standard isolation and culture techniques (see the experimental procedures), we isolated and cultured tissue-derived fibroblasts that were RFP^+^ or stained positive for αSMA from freshly harvested ears, tail tips, kidneys, lungs, fetal tissue (E13.5 and E18), orthotopically implanted 4T1 mouse mammary tumors, and spontaneously arising mammary tumors from the MMTV-PyMT mouse strain [33] (**Table 1, Fig. S1**). To ascertain the putative resiliency of fibroblasts, we generated an extensive tissue stress environment by allowing tissue to decay at room temperature post-mortem. To this end, mice were euthanized (see the experimental procedures), and the bodies were left at room temperature in a hydrated cloth to allow for decay for 12, 24, 48, 72, 96, 120, and 168 hours (**Table 1**).

**Table 1:** Fibroblast culture from different tissues. Table showing successful ex vivo expansion of cells from post-mortem, decaying tissues. Green check 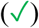 and red cross (X) denotes successful and unsuccessful ex vivo expansion monitored for 14 days post initiation of culture, respectively. Numbers indicate how many successful isolations of cells out of total number of mice used for isolations. ^#^ For example, ‘2/3’ indicate two successful expansions of cells from the kidneys of 3 distinct mice that were allowed to decay for 12-hours prior to tissue harvest.

| Tissue | 0 hr | 12 hr | 24 hr | 48 hr | 72 hr | 96 hr | 120 hr | 168 hr |
| --- | --- | --- | --- | --- | --- | --- | --- | --- |
| Ear | ✓<br>9/9 | ✓<br>9/9 | ✓<br>9/9 | ✓<br>11/11 | ✓<br>2/2 | ✓<br>2/2 | ✓<br>3/3 | X<br>0/1 |
| Tail | ✓<br>4/4 | ✓<br>4/4 | ✓<br>4/4 | ✓ / X<br>2/4 |  |  |  |  |
| Kidney | ✓<br>3/3 | ✓<br>2/3<sup>#</sup> | X<br>0/3 | X<br>0/3 |  |  |  |  |
| Lung | ✓<br>21/21 | ✓<br>26/26 | ✓ / X<br>1/7 | ✓ / X<br>1/7 |  |  |  |  |
| Fetus (E13.5 and E18) | ✓<br>2/2 | ✓<br>2/2 | ✓<br>2/2 | X<br>0/2 | X<br>0/1 |  |  |  |
| Tumor (4T1, n=1 and MMTV-PyMT, n=5) | ✓<br>6/6 | ✓<br>4/6 | X<br>0/2 | X<br>0/1 |  |  |  |  |

Culture experiments demonstrated that fibroblasts were successfully expanded ex vivo from tissues even after they were allowed to decay for 12-hours at room temperature (**Table 1, Fig. S1)**. Spindle-shaped fibroblasts were isolated and cultured from all tissues tested in the 12-hours post-mortem mice and were characterized by expression of αSMA, either by detection of αSMA-RFP signal or immunostaining of αSMA (**Fig. S1**). After 24-hours, fibroblasts could be isolated and cultured from the ear, tail tip, and fetus, but not from mammary tumors, kidneys, and lungs (**Table 1, Fig. S1**). After 48-hours, only ear and tail tips were amenable for ex vivo expansion of fibroblasts (**Table 1, Fig. S1**). From 48-hours and up until 120-hours, fibroblasts could still be cultured from ear of mice that were highly decomposed (**Table 1, Fig. S1**). Our results show fibroblasts from that distinct tissue beds show different resilience to tissue decay based on their successful ex vivo expansion. Interestingly, carcinoma associated fibroblasts do not present with a superior growth or survival advantage compared to non-neoplastic tissues in our experimental setting.

Next, we evaluated whether cells expressing epithelial or mesenchymal markers differentially survive in post-mortem decaying tissue. We performed flow cytometry analysis of cells obtained from digested ear and lungs αSMA-RFP mice and immunolabeled for the epithelial marker E-Cadherin (**Fig. S2 and S3**). Approximately 10% of cells were E-Cadherin^+^ in both ears and lungs, in contrast with 8% of cells in the ear and 3% of cells in the lung were αSMA-RFP^+^ (**Fig. S2 and S3**). Cells positive for both E-Cadherin and αSMA-RFP were rare in both ears and lungs (less than 1%, data not shown). In 12-hours post-mortem ear tissue decaying at room temperature, the percentage of live E-Cadherin^+^ (**Fig. 1A**) and αSMA-RFP^+^ cells (**Fig. 1B**) was not significantly altered at 12-hours post-mortem. In contrast, the percentage of E-cadherin^+^ cells that were alive was significantly reduced in lungs (**Fig. 1C**, p<0.05) and the percentage of αSMA-RFP^+^ cells that remained alive was not significantly changed (**Fig. 1D**), suggesting that epithelial cells in the lung are more sensitive to organismal decay compared to epithelial cells in the ear. Despite continued presence of E-cadherin^+^ cells in the 12-hours post-mortem tissue, these cells were not amenable for culture, suggesting a diminished survival program in this experimental setting. Cells cultured from the ear of αSMA-RFP mice that were allowed to decay at room temperature for increasing amount of time were labeled for the fibroblasts-enriched, mesenchymal markers FSP1 (S100A4) and vimentin (**Fig. S4**). Nearly 100% of cells isolated at all time points (0-hour to 48-hours) of post-mortem tissue expressed FSP1 and vimentin (**Fig. 1E-F**), and 80% on average remained αSMA-RFP^+^ (**Fig. 1G**), supporting that majority of isolated cells are fibroblasts.

**Figure 1:**
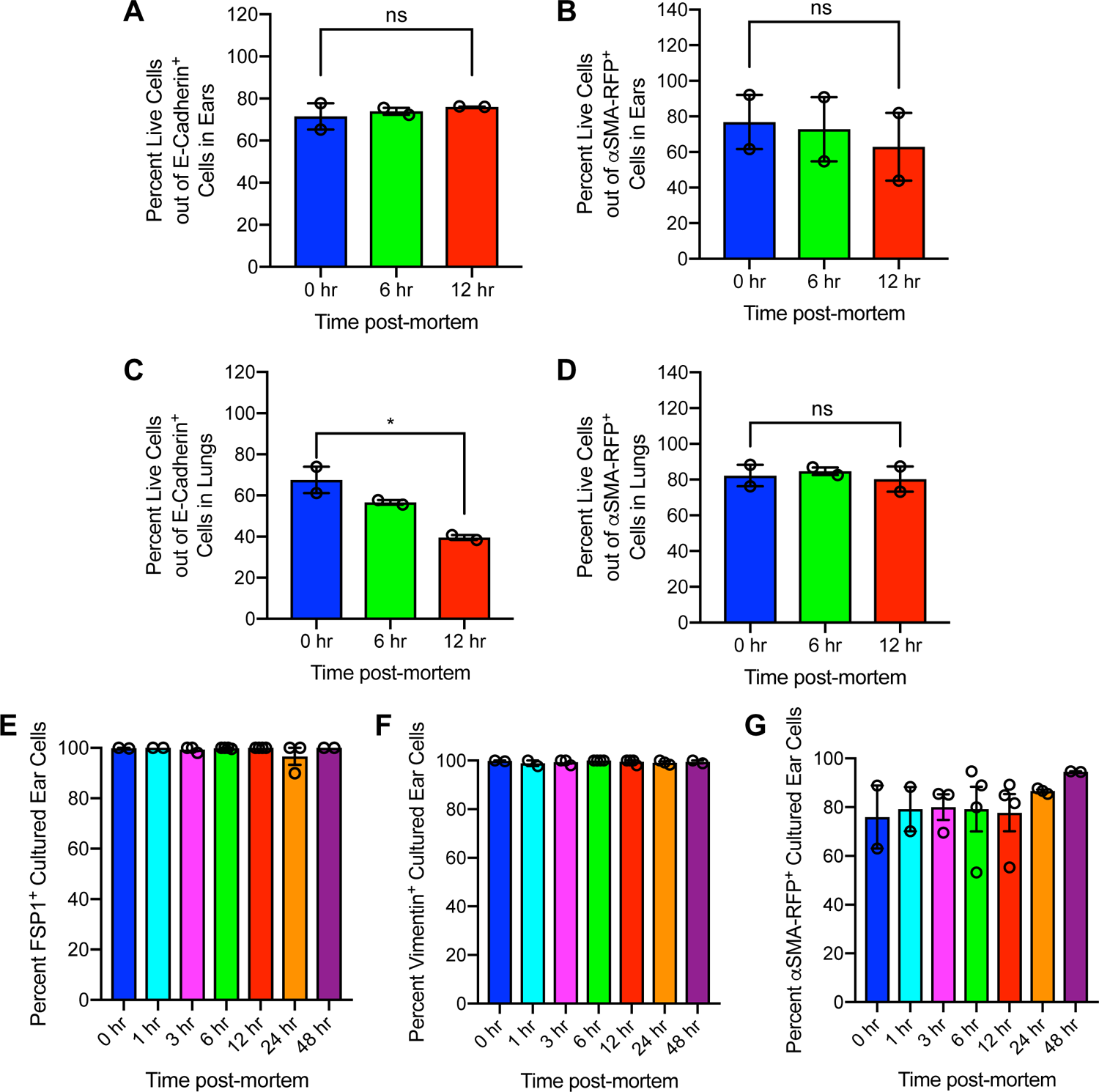
Fibroblasts survive post-mortem decay. (A and B) Flow cytometry analysis of live E-Cadherin^+^ (A) and αSMA-RFP^+^ (B) cells from digested ear (n=2 mice per group). (C and D) Flow cytometry analysis of live E-Cadherin^+^ (C) and αSMA-RFP^+^ (D) cells from digested lung (n=2 mice per group). One-way ANOVA was performed with Holm-Sidak’s multiple comparison post-hoc test comparing 0-hour post-mortem to all time points. * P<0.05. (E, F and G) Quantification of FSP1^+^ (E), vimentin^+^ (F) and αSMA-RFP^+^ (G) fibroblasts isolated and cultured for 14 days from ear harvested at the indicated post-mortem times (n=2-4 independent cell isolations).

We next sought to evaluate the putative mechanism(s) associated with fibroblasts resilience to severe post-mortem stress conditions in decaying tissue. Telomerase reverse transcriptase (TERT) has been implicated in cell survival in response to stress through both enzymatic activity-dependent [34] and independent [35] mechanisms. We isolated ear fibroblasts from third generation TERT deficient mice (G3^TERT-ER^) to determine whether loss of TERT would influence the survival of fibroblasts in decaying post-mortem tissue. Fibroblasts were successfully isolated and recovered from decaying post-mortem tissue up-until 96 hours after death of the mice, suggesting that TERT is dispensable for post-mortem survival in this experimental setting (**Fig. S5**).

Previous studies have demonstrated that senescent cells accumulate in damaged tissues [36]. To determine whether post-mortem fibroblasts displayed increased presence of markers suggestive of cellular senescence, senescence-associated β-galactosidase staining was performed (**Fig. S6A**). While insignificant differences in the percentages of β-galactosidase^+^ cells were observed in fibroblasts cultured after 24-hours post-mortem, an increase in their number was detected at 48-hours post-mortem (p<0.001, **Fig. S6B**). These results suggest that sustained survival of fibroblasts in post-mortem decaying tissue is associated with pathways connected to cellular senescence at later time points. Despite increased senescence, fibroblasts continue to survive until 120-hours post-mortem (**Table 1, Fig. S1)**.

### Gene expression profiling suggests post-mortem lung fibroblasts activate autophagy and lysosomal pathways

RNA-seq analysis was performed to identify transcriptional changes between fresh (0-hour) and 12-hours decayed post-mortem ear and lung fibroblasts. Both ear- and lung-derived fibroblasts expressed mesenchymal genes and lacked expression of epithelial genes (**Fig. 2A-B**), supporting that the cells isolated in this experimental setting are primarily fibroblasts (**Fig. 1E-G**). The isolated fibroblasts also expressed type I collagen (*Col1a1*) without any change in its expression between 0-hour and 12-hours post-mortem (**Fig. 2A-B**). Relatively high levels of expression of FSP1 (*S100a4*) and Vimentin (*Vim*) was also noted, agreeing with our immunolabeling studies (**Fig. 1E-F**). Supervised clustering revealed samples cluster with their tissue of origin (lung or ear) and fibroblasts isolated at 12-hours post-mortem had distinct transcriptional profiles compared to fibroblasts isolated at 0-hour post-mortem (fresh) (**Fig. 2C**). Volcano plots identified significantly differentially expressed genes in post-mortem ear and lung fibroblasts (**Fig. 2D, Fig. S7A-D**). Gene set enrichment analysis (GSEA) indicated upregulation of pathways associated with cardiomyopathy, cytokine interactions, and JAK/STAT signaling in 12-hour post-mortem ear fibroblasts (**Fig. 2E**). The cardiomyopathy pathway captured by GSEA possibly reflects post-mortem fibroblasts presenting with an enhanced cytoskeletal contractile apparatus with actin and myosin related genes (*Myh4, Xirp1, Mybpc1, Myl1*) and associated metabolism genes (*Ckmt2, Ppp1r3a*) (**Fig. 2D, Fig. S7A**). In contrast, DNA repair, cell cycle, metabolism, and ribosome pathways were downregulated in 12-hours post-mortem ear fibroblasts (**Fig. 2E**). In lung post-mortem fibroblasts, genes implicated in cell adhesion (*Dnah2, Ctnna2, Fbln1*), and pathways associated with ribosome biogenesis, replication, spliceosome formation, and DNA repair were downregulated (**Fig. 2F**). Post-mortem fibroblasts upregulate genes associated with lectin (*Clec12a, Cd72*), cytokine (*Ccl12*), and calcium signaling (*Trpc6*), and gene expression pathways indicate positive enrichment for cytokine receptor interaction, lysosome (*Enpep*), and regulation of autophagy (*Nup107*) pathways (**Fig. S7E**).

**Figure 2:**
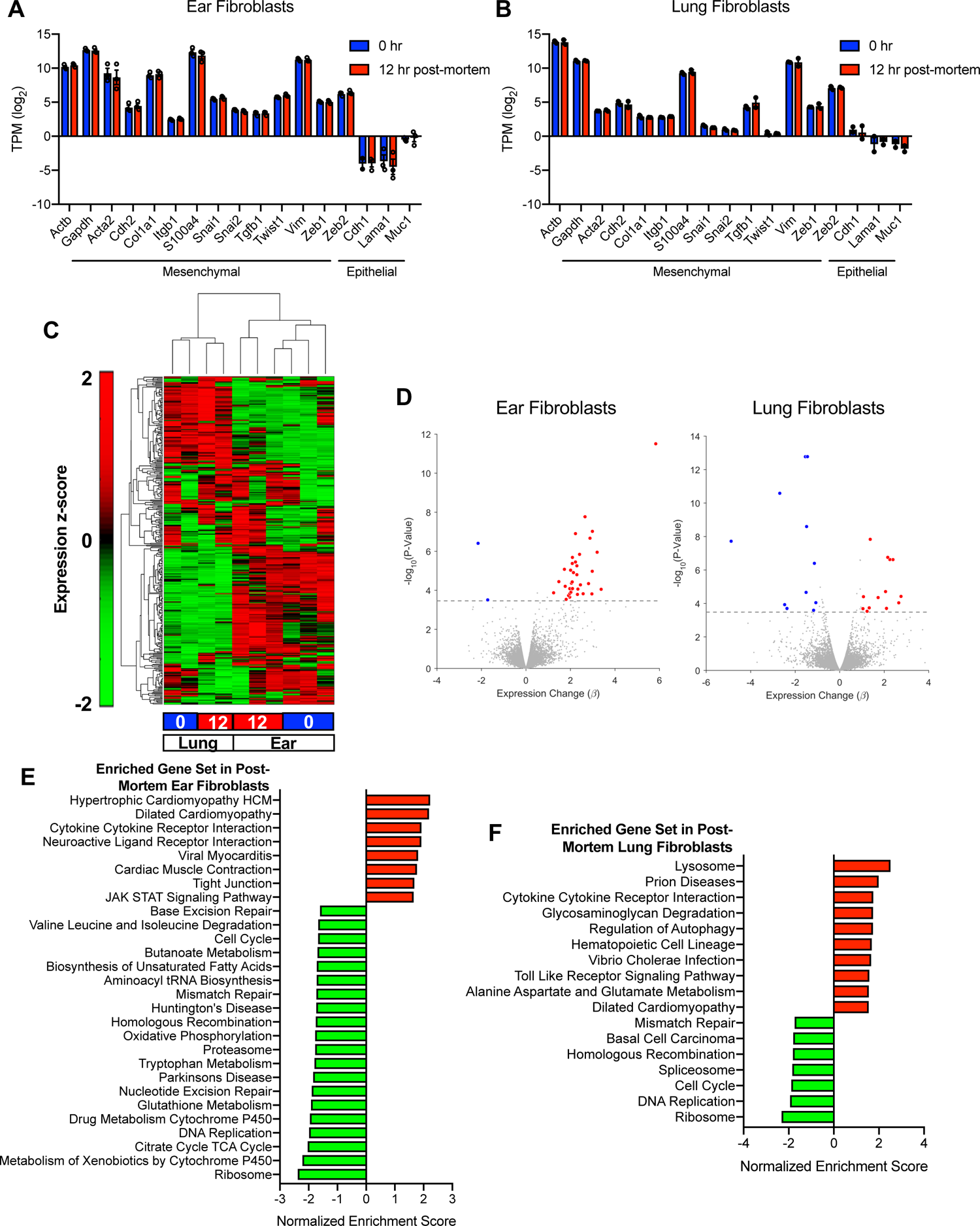
Gene expression profiling of fibroblasts isolated from post-mortem decaying tissue. (A and B) mRNA expression quantified by RNA-sequencing (transcripts per million, TPM) of 0-hour (blue) and 12-hours (red) post-mortem ear (A) and lung (B) fibroblasts compared to housekeeping genes (*Actb* and *Gapdh*). (C) Heat map of supervised clustering of mRNA expression in lung (n=2 independent fibroblast isolations) and ear fibroblasts (n=3 independent fibroblast isolations). Red indicates higher expressed genes and green indicates lower expressed genes. (D) Volcano plots of significantly differentially regulated genes (|β|>1, q-value<0.1) in 12-hours compared to 0-hour post-mortem fibroblasts (downregulated genes: blue, upregulated genes: red, genes that are not significantly differentially regulated: grey). Dashed line indicates q-value cutoff. (E and F) Pathways identified as significantly enriched (q-value<0.1) by gene set enrichment analysis (GSEA) in ear (E) and lung (F) fibroblasts isolated at 12-hours compared to 0-hour post-mortem.

To confirm whether autophagy is indeed upregulated in lung fibroblasts isolated from post-mortem decaying lung tissue, we profiled the modification/conversion of light chain 3B-I (LC3B-I) protein into LC3B-II protein (**Fig. 3A, Fig. S8**), indicative of autophagosome formation. Increased expression of LC3B-II was present in 12-hours post-mortem fibroblasts compared to freshly isolated 0-hour post-mortem fibroblasts (**Fig. 3B**). Next, immunostaining was performed to characterize the presence of autophagosomes (LC3B^+^ foci), lysosomes (lysosomal-associated protein 1, LAMP-1^+^ foci) and autolysosomes (LC3B^+^ LAMP-1^+^ foci) (**Fig. 3C**). Image quantification revealed that the number of autophagosomes and lysosomes per cell was insignificantly altered, while autolysosomes per cell were significantly increased in 12-hours post-mortem fibroblasts when compared with freshly isolated 0-hour post-mortem fibroblasts (**Fig. 3D-F**, p<0.01).

**Figure 3:**
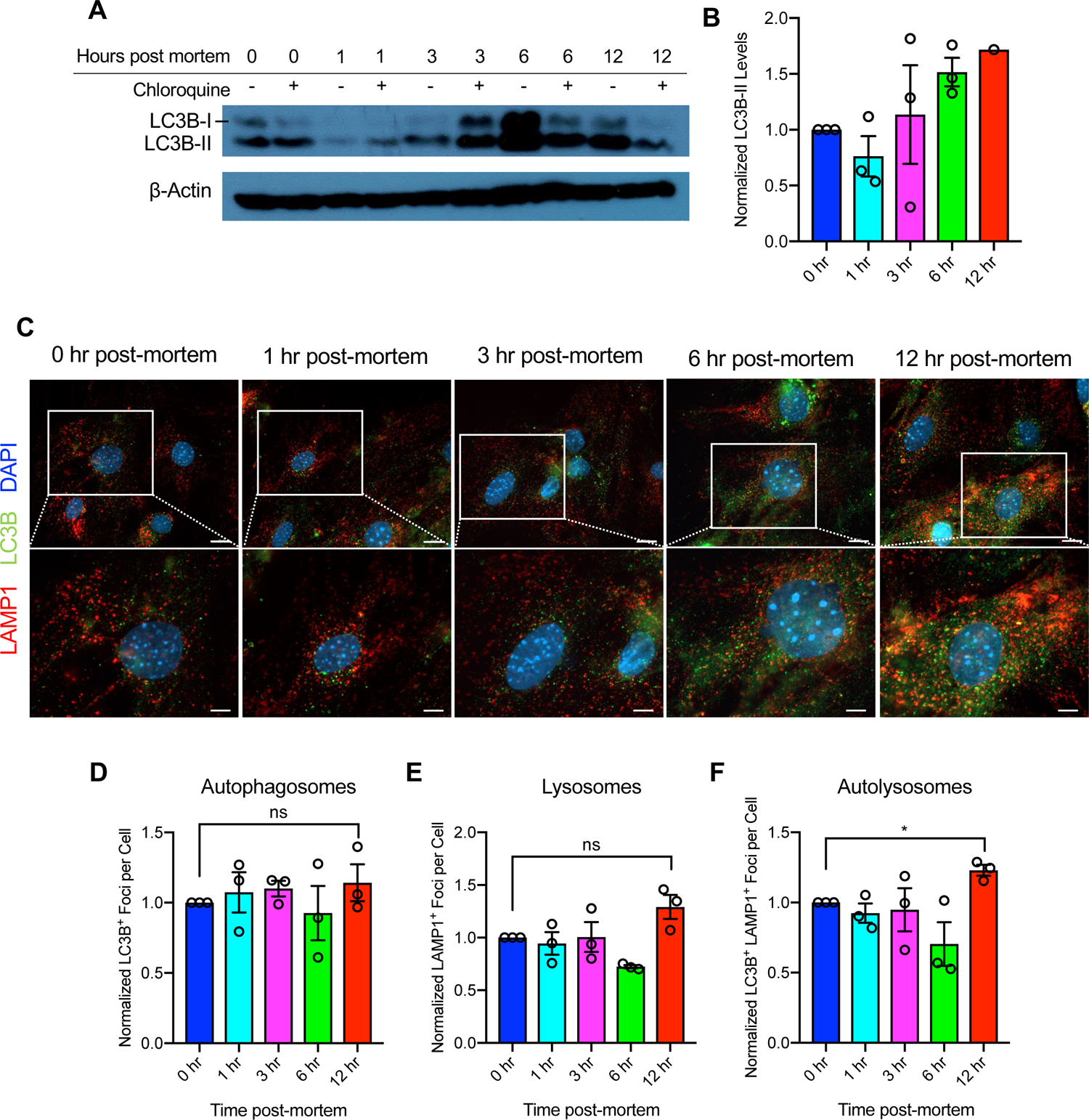
Augmented autophagy in fibroblasts isolated from post-mortem decaying tissue. (A) Western blot analysis of LC3BI/II in lung fibroblasts treated with chloroquine. Loading control: β-Actin. (B) Quantification of LC3BII protein levels. Expression levels are normalized to β-Actin levels and to 0-hour postmortem fibroblasts. Quantification performed based on blots in Fig. S8. (C) Representative images of lung fibroblasts immunostained for LAMP1 (red), LC3B (green), and nuclei (blue). Lower panel: digital zoom of boxed area in upper panel. Scale bar: 20 µm for full size images, 10 µm for zoomed images. (D, E, and F) Quantification of LC3B^+^ (D), LAMP1^+^ (E), and LC3B^+^LAMP1^+^ (F) foci per cell (n=3 independent cell isolations per group). Data is normalized to 0 hr fibroblasts and one-sample t test was performed comparing each timepoint to a theoretical mean of 1. * P<0.05.

### Increased number of exosomes traffic to lysosomal compartments in post-mortem fibroblasts

To determine whether endocytosed extracellular vesicles traffic via lysosomes in 12-hours post-mortem fibroblasts differently when compared with 0-hour post-mortem fibroblasts, we treated lung fibroblasts with fluorescently labeled 4T1 mammary cancer cell-derived exosomes and immunolabeled recipient cells for LC3B and LAMP-1 to evaluate co-localization of exosomes with autophagosomes, lysosomes and/or autolysosomes (**Fig. 4A**). Flow cytometry (FACS) analysis of exosome-incubated fibroblasts revealed an insignificant difference in the level of exosome uptake (endocytosis) in 12-hours post-mortem fibroblasts when compared with freshly isolated 0-hour post-mortem fibroblasts (**Fig. S9A-B**). Image analysis showed increased co-localization of fluorescently labeled exosomes with lysosomes but not with autophagosomes and autolysosomes in the 12-hours post-mortem fibroblasts when compared to the 0-hour post-mortem fibroblasts (**Fig. 4B-D**). Together, the data indicate that while exosomes entry into fibroblasts is not altered, exosomes are increasingly targeted to degradative lysosomes in 12-hours post-mortem fibroblasts.

**Figure 4:**
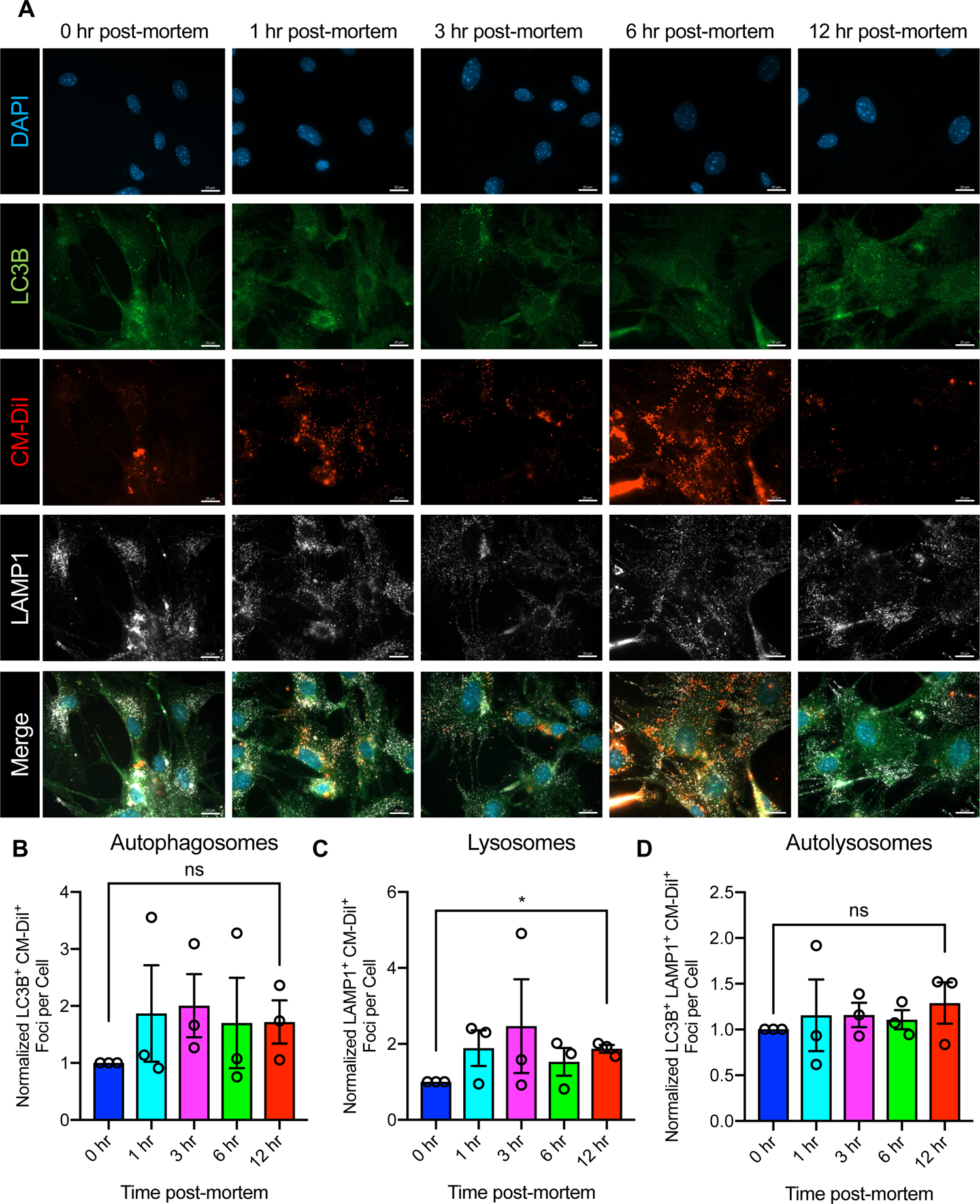
Altered exosome trafficking in post-mortem fibroblasts. (A) Representative images of lung fibroblasts treated with CM-DiI labeled 4T1 exosomes (red) and immunostained for LC3B (green), LAMP1 (white) and nuclei (blue). Scale bar: 20 µm. (B, C, and D) Quantification of LC3B^+^CM-DiI^+^ (B), LAMP1^+^CM-DiI^+^ (C), and LC3B^+^LAMP1^+^CM-DiI^+^ (D) foci per cell (n=3 independent cell isolations per group). Data is normalized to 0 hr fibroblasts and one-sample t test was performed comparing each timepoint to a theoretical mean of 1. * P<0.05.

## Discussion

This study shows a dramatic capacity for survival and resilience of fibroblast in decaying post-mortem mouse tissue. Cells expanded ex vivo from decaying tissues showed mesenchymal markers expression, suggesting a unique property of fibroblasts to survive severe stress condition and autolytic decay of tissue for up to 120-hours. This feature of fibroblasts is likely what enabled their culture and evaluation from post-mortem human tissue [37]. Preservation of cells from an unlikely/sudden death of an individual can have profound implication for tissue matched therapies for relatives and others in need, including generation of iPSCs.

Our studies show that fibroblasts can be cultured from decayed post-mortem murine ears even after 120 hours. This was not observed with respect to internal organ tissues, which preserved viable fibroblasts until 12 hours post-mortem. The reason for this difference could be from the inherent heterogeneity of resident fibroblasts in distinct tissue. These differences may also reflect the nature of internal tissues, like lungs and kidneys, to undergo more rigorous autolysis from room-temperature decay when compared to tissue such as ears and tails (dermal tissue), and this needs further testing. Ear fibroblasts also showed enhanced resilience and capacity for ex vivo expansion post-mortem when compared to tail tip fibroblasts, suggesting that dermal fibroblasts from these distinct sources may also present with heterogeneity with implication for survivability post-mortem.

One reason for fibroblasts to survive post-mortem tissue decay could also be due specific epigenetic modifications. Although cancer associated fibroblasts undergo specific epigenetic alteration that likely aid in their sustained proliferation and survival [22], they do not seem to culture better than normal fibroblasts in this study. More research needs to be conducted to understand the precise mechanism that governs the resiliency of normal fibroblasts.

Our study demonstrates an association of fibroblasts cultured from post-mortem decaying tissue with increased lysosomes and enhanced autophagy, when compared with freshly isolated tissue fibroblasts. Such difference is preserved despite culture adaptation. It remains to be evaluated whether survival of fibroblasts in decaying post-mortem tissue is due to exclusively intrinsic autophagy or due to lysosomal degradation of extracellular protein. While more experiments are required, in a proof-of-concept experiment, we used exosomes (a type of extracellular vesicles) to explore the potential of fibroblasts isolated from post-mortem tissue for endocytosis/fusion and uptake of exosomes. We show that this property remains intact in post-mortem fibroblasts. In contrast, compared to freshly isolated fibroblasts, the post-mortem fibroblasts adapt to trafficking the internalized exosomes to lysosomes, possibly for degradation of proteins to fuel biomass requirements.

Collectively, this study demonstrates the resiliency of normal fibroblast and further mechanistic study of this phenomenon will likely provide insight into cellular senescence, aging, and survival of environmental stress.

### Experimental Procedures

#### Animal studies

Mice were housed in individually ventilated cages with a 12-hours light/dark cycle at 21-23°C and 40-60% humidity and were fed standard lab chow. 8-12-week-old male and female mice were used for cell isolation, except in the case of fetal and tumor tissue isolations where only female mice were utilized. Mice were euthanized via CO_2_ asphyxiation at a displacement rate of 20% per minute. After chest rising had ceased, mice were cervically dislocated and the chest cavity opened. *Acta2*-DsRed (αSMA-RFP) mice were generated in our laboratory and previously described [31]. Third generation TERT-ER (G3^TERT-ER^) mice were kindly provided by Dr. Ronald DePinho and previously described [38]. MMTV-PyMT mice were purchased from Jackson Laboratory (FVB/N-Tg (MMTV-PyVT)634Mul/J) and maintained on Balb/C background. For experiments analyzing light chain 3B (LC3B) and lysosomal-associated membrane protein 1 (LAMP-1) expression, wild-type BALB/c mice (Charles River) were used. αSMA-RFP and G3^TERT-ER^ mice were maintained on BALB/c and C57BL/6 genetic backgrounds, respectively. For isolation from fetal tissue, the pregnancy of a wild-type C57 female was confirmed via presence of vaginal plug (denoted as E0.5). Dams were euthanized at E13.5 and E18 for cell isolation. For 4T1 mammary tumor experiments, mice were orthotopically injected with 1×10^6^ 4T1 cells in 1X PBS and tumors grown to ∼1500 mm^3^. All manipulations were performed with MD Anderson Cancer Center IACUC approval. After sacrifice, tissue was collected within minutes (fresh tissue, 0-hour post-mortem), then mice were covered in PBS-soaked gauze and stored at room temperature for the indicated times post-mortem.

#### Cell isolation and culture

For fibroblast isolation, similar sized pieces of tissues were collected (approximately 1 cm), minced into ∼1 mm^3^ pieces and digested in 400 U/mL collagenase IV (Gibco) in RPMI (Corning) with 1% penicillin-streptomycin with antimycotics (PSA, Corning) at 37°C overnight. To isolate from fetal tissue, the placenta was removed as well as the head of the embryo prior to digestion as described above. Tumor tissue was collected from the tumor periphery to avoid necrotic areas of one 4T1 tumor-bearing mouse and 5 MMTV-PyMT mice. Media was replaced the following day to RPMI with 10% FBS (Gemini) with 1% PSA. All fibroblast analyses were performed at passage two (approximately 14 days in culture) to avoid any artifacts of cell culture-associated senescence. In addition, cells were isolated from 3 independent mice for each experiment, unless otherwise noted. 4T1 cells were obtained from the MD Anderson Cancer Center Characterized Cell Line Facility and cultured in DMEM (Corning) with 10% FBS and 1% penicillin-streptomycin (Corning). 4T1 cells were STR validated and tested for rodent pathogens and mycoplasma.

#### FACS analysis of cells from lungs

Lungs from αSMA-RFP mice were minced into ∼1 mm^3^ pieces and digested in 300 U/mL collagenase I (Gibco) in DMEM (Corning) for 1 hour at 37°C with shaking. Digested tissue was passed through an 18G needle followed by 21G needle prior to filtering through a 40 µm cell strainer. Samples were incubated with 20 µg/mL DNAse I (Sigma Aldrich) for 5 minutes on ice, followed by centrifugation and treatment with ACK lysis buffer (Gibco) for 10 minutes at room temperature. Samples were blocked with 5 µg/mL CD16/32 antibody (BD #553141) in FACS buffer (PBS with 2% FBS) for 20 minutes, then stained with anti-E-cadherin eFluor 660 (Invitrogen #50-3249-82, 1:100) or corresponding isotype antibody (Invitrogen #50-4301-80, 1:100) for 30 minutes on ice. After antibody staining, cells were stained with Fixable Viability Stain 510 (BD Biosciences) and analyzed with a BD LSR Fortessa X-20 flow cytometer. Data was analyzed using FlowJo software (TreeStar Inc).

#### Immunocytochemistry

For FSP1 and vimentin staining, cells were fixed in 4% paraformaldehyde for 10 minutes at room temperature. Cells were permeabilized with 1% Triton-X for 15 minutes at room temperature followed by blocking with 2% BSA at room temperature for 1 hour. Cells were stained with anti-FSP1/S100A4 (Dako #5114, 1:100) for 1 hour at room temperature followed by staining with anti-rabbit Alexa Fluor 488 (Invitrogen #A11008, 1:1000) and anti-vimentin Alexa Fluor 647 (Cell Signaling #9856S, 1:100) for 1 hour at room temperature. For embryonic and tumor fibroblasts, cells were stained with anti-αSMA Cy3 (Sigma Aldrich #C6198, 1:100) for 1 hour at room temperature. Coverslips were mounted and sealed with Fluoshield with DAPI (Sigma Aldrich). Images were acquired at 20x magnification on a Zeiss Axio Observer Z1 inverted fluorescent microscope and positive cells quantified in ImageJ (NIH). Biological replicates for each time point post-mortem consisted of 3 technical replicates with 8 images per technical replicate. Senescence was analyzed using Senescence β-Galactosidase Staining Kit (Cell Signaling #9860) according to manufacturer’s instructions. Nuclei were stained with Hoechst 33342 (Invitrogen #H3570) and cells imaged at 10x magnification with a Zeiss Axio Observer Z1 inverted fluorescent microscope with 3 technical replicates of 8 images each per biological replicate. Counting of β-Galactosidase^+^ cells was performed in ImageJ. For LC3B and LAMP-1 staining, cells were fixed in ice-cold methanol for 10 minutes. Cells were permeabilized in 0.3% Tween-20 for 15 minutes and blocked in 2% BSA for 1 hour at room temperature. Cells were incubated with anti-LC3B (Cell Signaling #2775S, 1:100) and anti-LAMP-1/CD107a (Invitrogen #14-1071-82, 1:100) for 1 hour at room temperature followed by staining with anti-rabbit Alexa Fluor 488 (Invitrogen #A11008, 1:1000) and anti-rat Alexa Fluor 647 (Invitrogen #A21247, 1:500). Nuclei were stained with Hoechst 33342 and coverslips mounted with Vectashield Antifade Mounting Medium (Vector Labs). Slides were imaged at 63x magnification on a Zeiss Axio Observer Z1 Inverted fluorescent microscope. Biological replicates for each time point post-mortem consisted of 3 technical replicates with 8 images per technical replicate. Quantitative image analysis was performed in Matlab 2016a using custom written scripts. In brief, nuclei were segmented by thresholding using Otsu’s method. For extracting puncta such as vesicles or exosomes, a bandpass filter was applied before thresholding. Only puncta larger than 0.26 µm^2^ were quantified. Likewise, for co-localization analysis two puncta from the two separate channels also had to overlap by at least 0.26 µm^2^. For each image, puncta number was normalized to total cell area.

#### Western blot

Protein was isolated from cells in 8 M urea with 2.5% SDS and complete protease inhibitor cocktail (Roche) and quantified by BCA assay. Protein (20 µg) was separated electrophoretically on a polyacrylamide gel and transferred to a PVDF membrane using a semi-dry transfer system (BioRad). Membranes were blocked for 1 hour with 5% BSA and incubated overnight at 4°C with LC3B antibody (Cell Signaling #2775, 1:1000). Membranes were washed with TBS-T three times then incubated with HRP-conjugated anti-rabbit secondary antibody (Abcam #ab16284, 1:1000) for 1 hour at room temperature. Membranes were washed with TBS-T three times and developed using West Pico ECL reagent (GenDepot). Blots were then washed with PBS and incubated with HRP-conjugated anti-β-actin (Sigma Aldrich, 1:40,000) for 1 hour, washed three times with TBS-T, and developed as described above. Blots were quantified in ImageJ comparing LC3B-II levels to β-actin and then normalizing to 0 hours post-mortem.

#### Exosome isolation

For exosome collection, 4T1 cells were cultured in normal growth media in T225 flasks until 70% confluency. Flasks were washed with PBS and incubated in serum-free DMEM for 48 hours. Media was collected and sequentially centrifuged at 800 x g for 5 minutes followed by 2,000 x g for 10 minutes. Media was filtered with a 0.2 µm filter and centrifuged at 100,000 x g for 2 hours at 4°C in a SW 32 Ti rotor (Beckman Coulter). Exosome pellets were resuspended in 100 µL 1X PBS and exosome number quantified by Nanosight analysis (Malvern Nanosight LM10).

#### Exosome labeling and treatment of cells

4T1 exosomes were labeled by incubating approximately 2.4×10^12^ exosomes with 1 µg/mL CM-DiI (Invitrogen) in 1X PBS for 5 minutes at 37°C followed by a 15-minute incubation at 4°C. Samples were washed in PBS in a SW 41 Ti rotor (Beckman Coulter) at 200,000 x g for 2.5 hours at 4°C, followed by resuspension in RPMI with 10% exosome-depleted FBS and 1% PSA for treatment of cells. For exosome treatment of lung fibroblasts, 4×10^4^ cells were seeded in 24 well plates. The following day, cells were treated with labeled exosomes (approximately 2×10^6^ exosomes/cell) for 24 hours prior to fixation with methanol for immunostaining (as described above) or trypsinization for analysis using a BD LSR Fortessa X-20. Flow cytometry data was analyzed using FlowJo software.

#### RNA-seq analysis

RNA was isolated using AllPrep DNA/RNA kit (Qiagen), quantified using Qubit HS RNA kit (Invitrogen) and quality assessed with High Sensitivity RNA ScreenTape analysis (Agilent). Sequencing was performed according to manufacturer’s instructions and analyzed on a HiSeq 4000 (Illumina) with 76 nucleotide paired-end format. Paired-end next generation sequencing reads were quantified using kallisto [39] (v0.44.0) with 100 bootstraps per sample. Differential gene expression was subsequently determined using sleuth [40] (v0.30.0) comparing 0-hour vs 12-hours for each site of fibroblast isolation. For hierarchical clustering, samples were clustered using genes with an average log_2_(fold change) of 0.75 or greater. Clustering was performed in Matlab 2016a using Spearman correlation as the distance metric. For volcano plots, significant genes were considered as those with a β>1 and false discovery rate (FDR) q-value < 0.1. Pathway analysis was performed using gene set enrichment analysis (GSEA) as previously described [41] using the pre-ranked function weighted by gene fold change and 1000 iterations with an FDR q-value < 0.1. Gene expression results (RNA sequencing data) have been deposited into GEO (GSE136708).

#### Statistical analysis

Statistical analysis was performed using one-way ANOVA and Holm-Sidak’s multiple comparison test in GraphPad Prism. For normalized data, a one-sample t test was performed. A p-value of less than 0.05 was considered significant.

## Acknowledgements

This study was primarily supported by the Cancer Prevention and Research Institute of Texas. D.J. McGrail was supported by Susan G. Komen PDF17483544 and NCI T32CA186892. L.B.G. was supported by the Norwegian Cancer Society. Other support came from NCI Grant CA016672(SMF) for the Sequencing and Microarray Facility and NCI Grant CA016672 for CCSG-funded Characterized Cell Line Core. We thank D. Lundy and L. Gibson for help with cell culture, Q. Ping, A.R. Haltom, and L.M. Becker for animal husbandry support, and M.L. Kirtley, Y. Wang, and S. Malasi for STR, pathogen, and mycoplasma testing.

---

**Figure S1: Fibroblasts can expand ex vivo from post-mortem decaying tissue**. Representative images of cells isolated from indicated tissues post-mortem. Ear, tail, kidney, and lung cells were isolated from αSMA-RFP mice at 0-48 hours. Images are phase contrast images with RFP image (red) overlaid, scale bar: 10 µm. For 72-120-hour ear cells, nuclei were stained with Hoechst (blue), scale bar: 100 µm. Fetal cells were isolated from a wild-type mouse and stained with αSMA (red) and Hoechst (blue), scale bar: 50 µm. Tumor cells are derived from a 4T1 mammary tumor and stained with αSMA (red) and Hoechst (blue), scale bar: 20 µm.

**Figure S2: FACS gating strategy and representative plots for ear tissue**. Cells were isolated from αSMA-RFP^+^ or wild-type BALB/c (negative control) mice. For FACS quantification, cells were gated based on SSC-A and FSC-A plot, followed by gating of singlets based on FSC-H and FSC-A plot. E-Cadherin^+^ and αSMA-RFP^+^ cells were gated based on unstained/negative control, respectively. Live E-Cadherin^+^ and αSMA-RFP^+^ cells were determined out of all E- Cadherin^+^ and αSMA-RFP^+^ cells, respectively.

**Figure S3: FACS gating strategy and representative plots for fibroblast isolated from post-mortem decaying lung tissue**. Cells were isolated from αSMA-RFP^+^ or wild-type BALB/c (negative control) mice. For FACS quantification, cells were gated based on SSC-A and FSC-A plot, followed by gating of singlets based on FSC-H and FSC-A plot. E-Cadherin^+^ and αSMA-RFP^+^ cells were gated based on unstained/negative control, respectively. Live E-Cadherin^+^ and αSMA-RFP^+^ cells were determined out of all E- Cadherin^+^ and αSMA-RFP^+^ cells, respectively.

**Figure S4**: **Post-mortem fibroblasts express mesenchymal markers**. Representative images of ear fibroblasts isolated from αSMA-RFP (red) mice at indicated times post-mortem and immunostained for FSP1 (green), vimentin (white), and nuclei (blue, Hoechst 33342). Images are individual channels of same visual field for each post-mortem time point. Scale bar: 50 µm.

**Figure S5**: **Survival of TERT-deficient post-mortem fibroblasts**. Brightfield images of ear fibroblasts isolated from third-generation telomerase deficient (G3 TERT^-/-^) mice at indicated times post-mortem. Lower panel: digital zoom of boxed area in upper panel. Scale bar: 100 µm for full size images, 50 µm for zoomed images.

**Figure S6**: **Enhanced senescence in post-mortem fibroblasts**. (A) Representative images of ear fibroblasts stained with Hoechst 33342 (blue) and for senescence-associated β-galactosidase (black). Lower panel: digital zoom of boxed area in upper panel. Scale bar: 100 µm for full size images, 50 µm for zoomed images. (B) Quantification of cells positive for β-galactosidase (n=2-4 independent cell isolations per group as indicated on plot). One-way ANOVA was performed with Holm-Sidak’s multiple comparison post-hoc test. *** P<0.001.

**Figure S7**: **Differential expression of genes in post-mortem fibroblasts**. (A and B) Top upregulated (A) and downregulated (B) genes in 12-hours compared to 0-hour post-mortem ear fibroblasts. (C and D) Top upregulated (C) and downregulated (D) genes in 12-hours compared to 0 hour post-mortem lung fibroblasts. (E) Enrichment plots for lysosome and regulation of autophagy pathways in lung fibroblasts.

**Figure S8**: **Uncropped Western blots of LC3B and β-Actin**.

**Figure S9**: **Exosome uptake by post-mortem fibroblasts**. (A and B) Flow cytometry analysis of lung fibroblasts treated with CM-DiI labeled 4T1 exosomes. Percent positive cells (A) and mean fluorescence intensity (MFI, B) of lung fibroblasts treated with CM-DiI labeled 4T1 exosomes. One-way ANOVA was performed with Holm-Sidak’s multiple comparison post-hoc test for (A) and one-sample t test was performed for (B). (C) Representative flow cytometry plots. Cont: untreated cells, + Exosomes: cells treated with labeled exosomes.

