## Supplementary Figures for "Autophagy is associated with survival and resiliency of fibroblasts in long-term decaying post-mortem parenchymal tissue"

### Figure S1

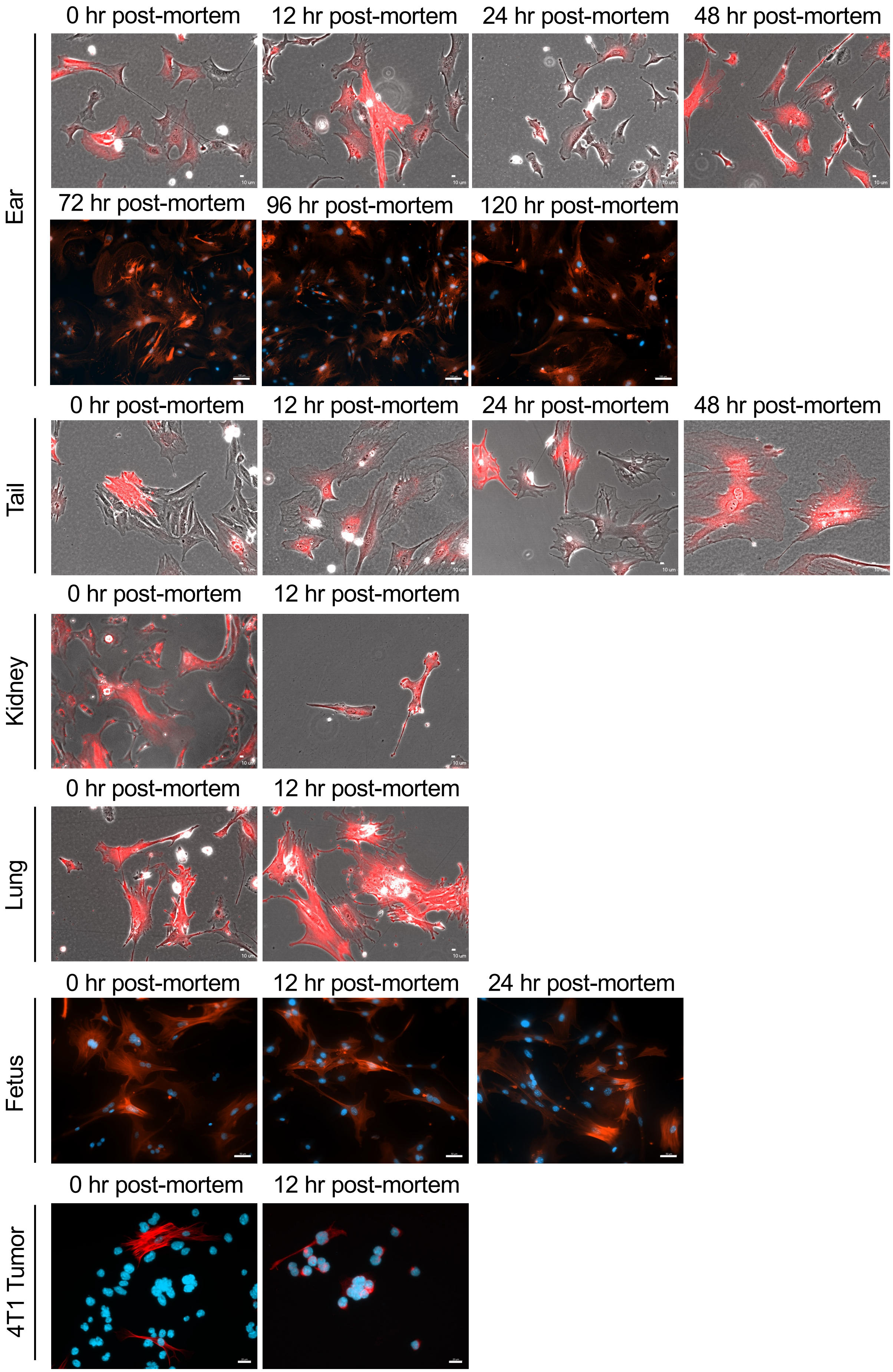
Ear
0 hr post-mortem
12 hr post-mortem
24 hr post-mortem
48 hr post-mortem
72 hr post-mortem
96 hr post-mortem
120 hr post-mortem
Tail
0 hr post-mortem
12 hr post-mortem
24 hr post-mortem
48 hr post-mortem
Kidney
0 hr post-mortem
12 hr post-mortem
Lung
0 hr post-mortem
12 hr post-mortem
Fetus
0 hr post-mortem
12 hr post-mortem
24 hr post-mortem
4T1 Tumor
0 hr post-mortem
12 hr post-mortem

### Figure S2

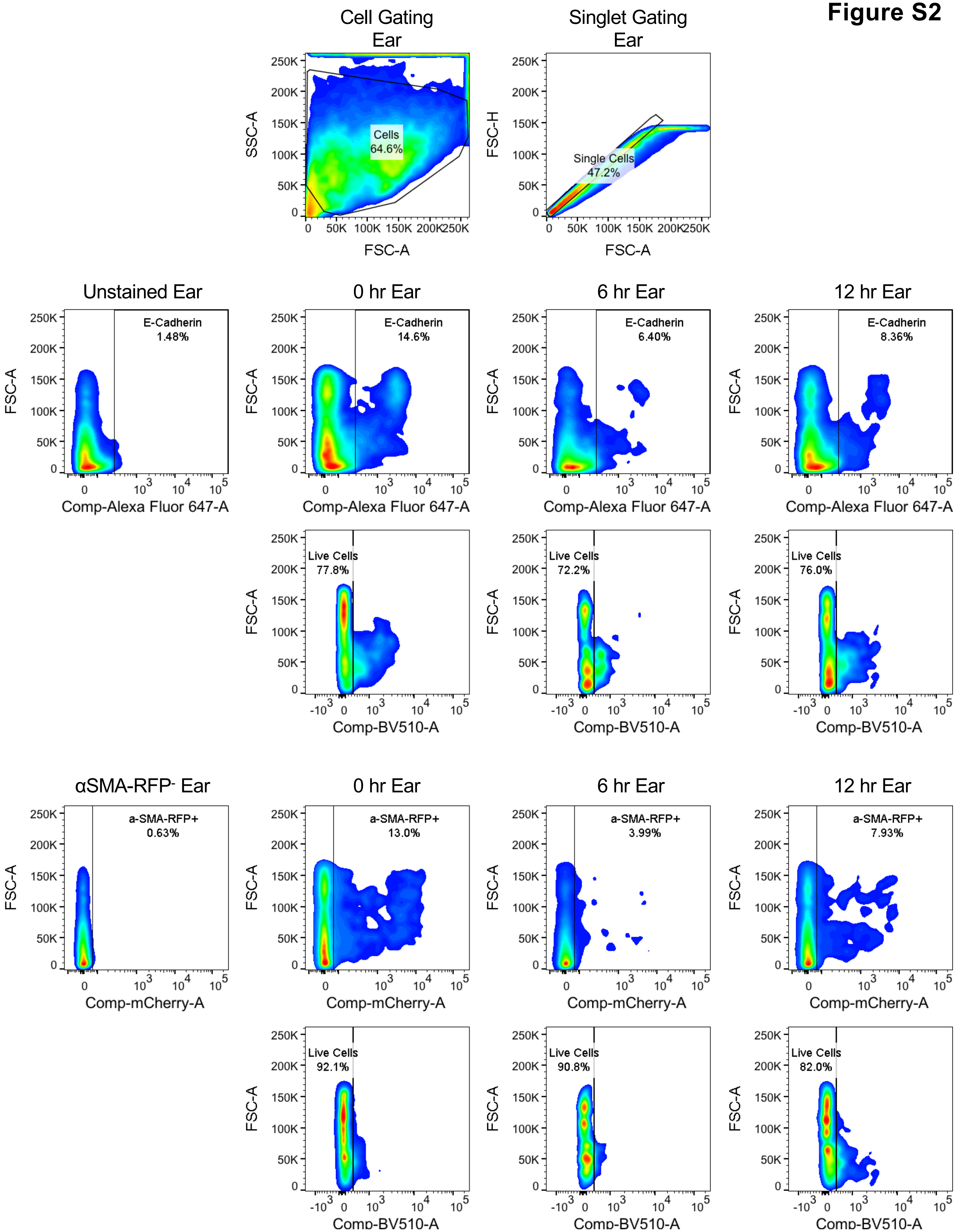

Cell Gating Ear
Cells 64.6%
SSC-A
FSC-A
Singlet Gating Ear
Single Cells 47.2%
FSC-H
FSC-A
Unstained Ear
0 hr Ear
6 hr Ear
12 hr Ear
E-Cadherin 1.48%
E-Cadherin 14.6%
E-Cadherin 6.40%
E-Cadherin 8.36%
FSC-A
Comp-Alexa Fluor 647-A
Live Cells 77.8%
Live Cells 72.2%
Live Cells 76.0%
Comp-BV510-A
αSMA-RFP- Ear
0 hr Ear
6 hr Ear
12 hr Ear
a-SMA-RFP+ 0.63%
a-SMA-RFP+ 13.0%
a-SMA-RFP+ 3.99%
a-SMA-RFP+ 7.93%
Comp-mCherry-A
Live Cells 92.1%
Live Cells 90.8%
Live Cells 82.0%
Comp-BV510-A

**Figure S3**

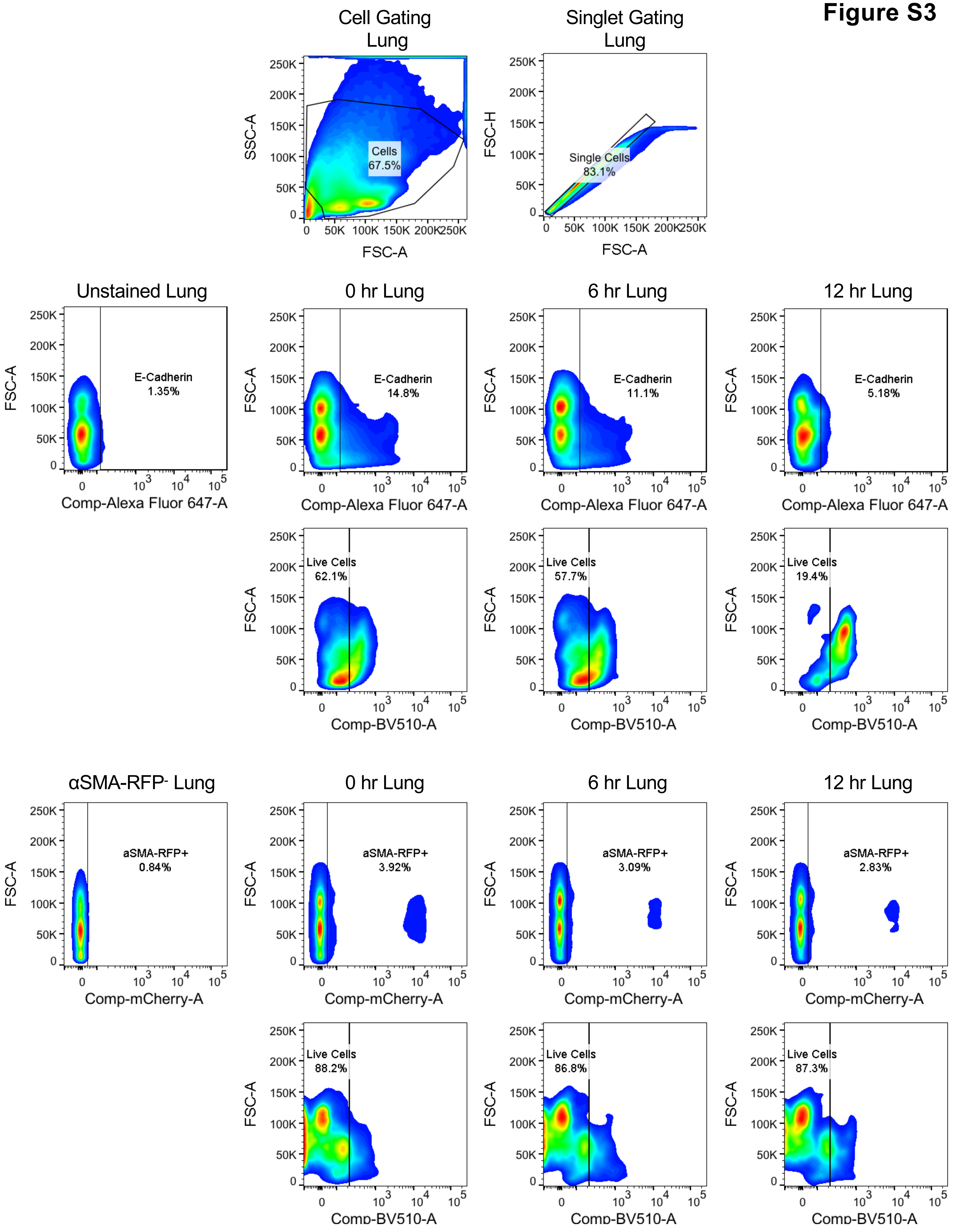
Cell Gating Lung
Cells 67.5%
SSC-A
FSC-A
Singlet Gating Lung
Single Cells 83.1%
FSC-H
FSC-A
Unstained Lung
E-Cadherin 1.35%
0 hr Lung
E-Cadherin 14.8%
6 hr Lung
E-Cadherin 11.1%
12 hr Lung
E-Cadherin 5.18%
Comp-Alexa Fluor 647-A
Live Cells 62.1%
Live Cells 57.7%
Live Cells 19.4%
Comp-BV510-A
αSMA-RFP- Lung
aSMA-RFP+ 0.84%
0 hr Lung
aSMA-RFP+ 3.92%
6 hr Lung
aSMA-RFP+ 3.09%
12 hr Lung
aSMA-RFP+ 2.83%
Comp-mCherry-A
Live Cells 88.2%
Live Cells 86.8%
Live Cells 87.3%
Comp-BV510-A
FSC-A

**Figure S4**

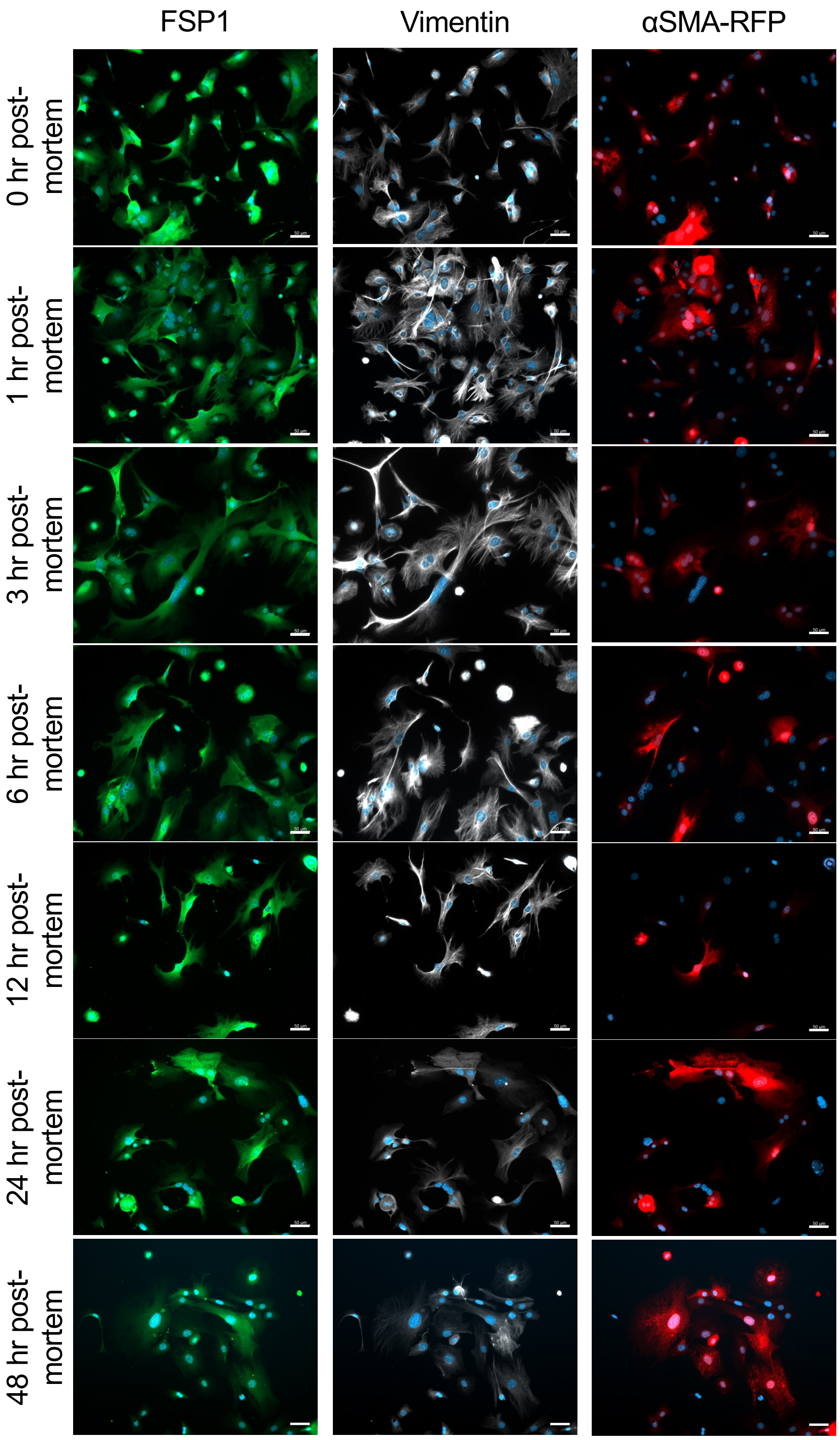
FSP1
Vimentin
αSMA-RFP
0 hr post-mortem
1 hr post-mortem
3 hr post-mortem
6 hr post-mortem
12 hr post-mortem
24 hr post-mortem
48 hr post-mortem

### Figure S5

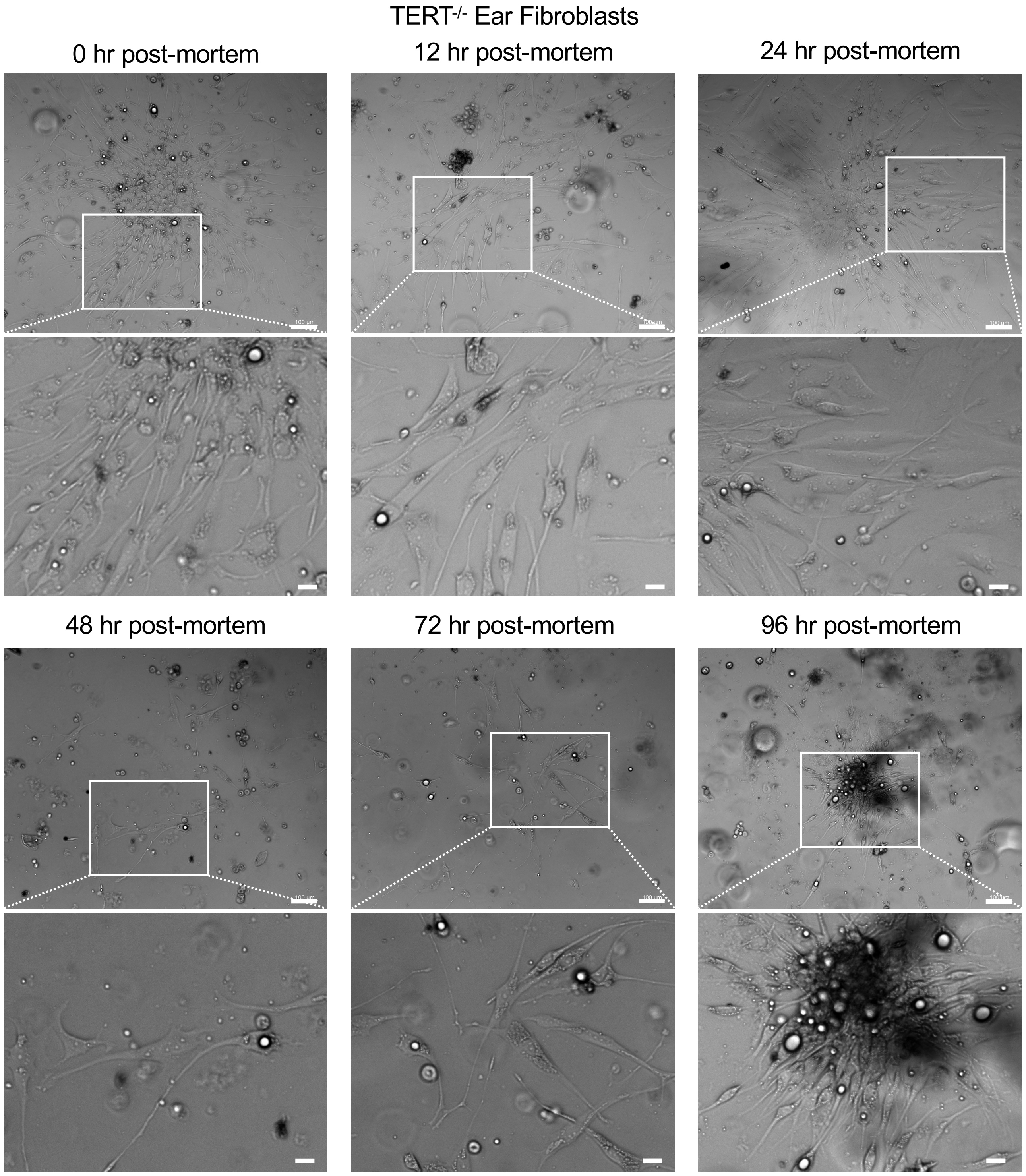
TERT-/- Ear Fibroblasts
0 hr post-mortem
12 hr post-mortem
24 hr post-mortem
48 hr post-mortem
72 hr post-mortem
96 hr post-mortem

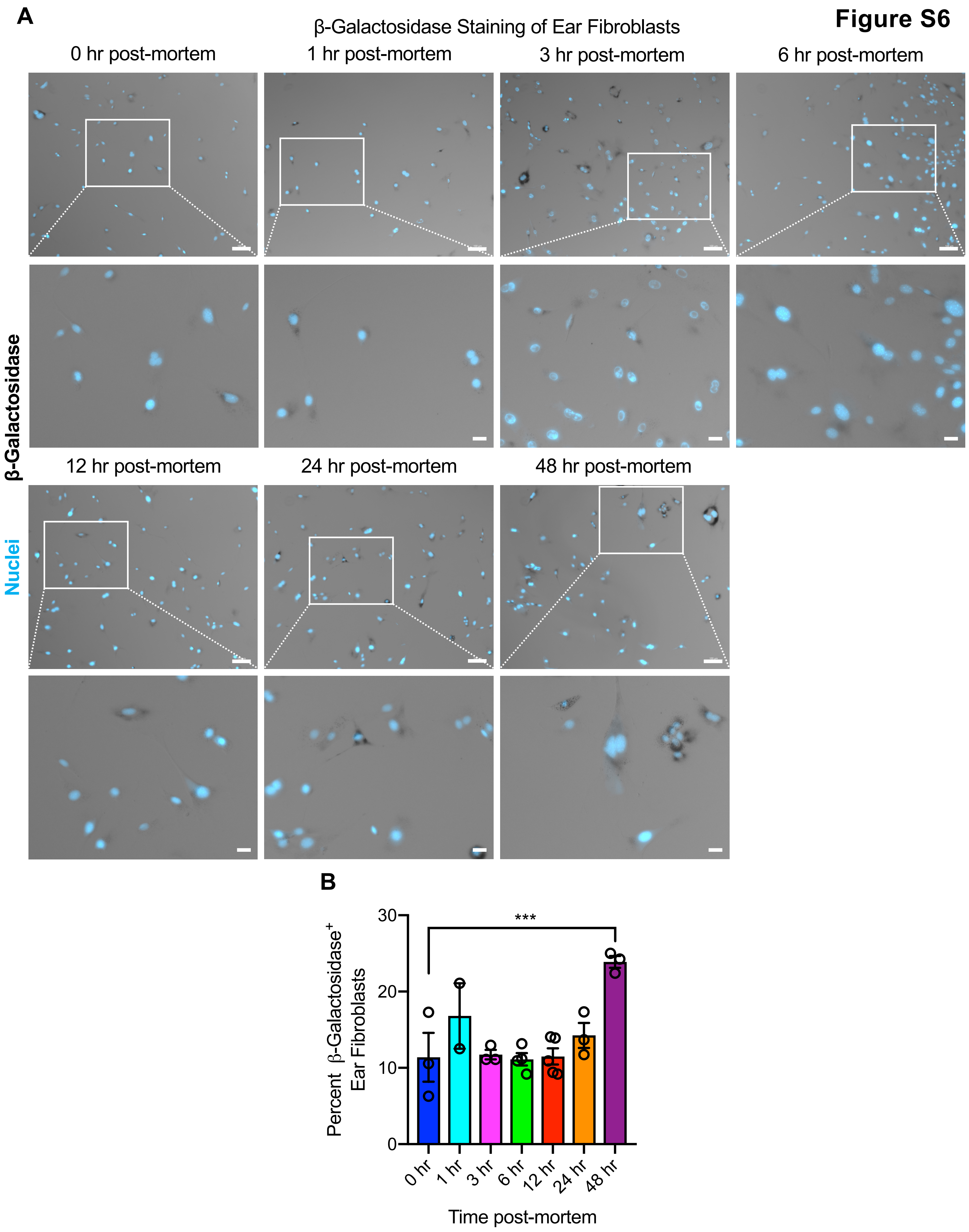
A
Figure S6
β-Galactosidase Staining of Ear Fibroblasts
0 hr post-mortem
1 hr post-mortem
3 hr post-mortem
6 hr post-mortem
β-Galactosidase
Nuclei
12 hr post-mortem
24 hr post-mortem
48 hr post-mortem
B
***
Percent β-Galactosidase+ Ear Fibroblasts
30
20
10
0
0 hr
1 hr
3 hr
6 hr
12 hr
24 hr
48 hr
Time post-mortem

### Figure S7

**A**

**Top Upregulated Genes in 12 Hour Ear Fibroblasts**

| Gene | Expression Change (β) | q-value |
|---|---|---|
| Txnl4a | 5.836 | 3.99E-08 |
| Myh4 | 3.379 | 4.04E-02 |
| Ckmt2 | 3.201 | 2.05E-03 |
| Chrnd | 2.988 | 3.99E-04 |
| Ppp1r3a | 2.982 | 9.03E-03 |
| Tceal7 | 2.965 | 5.75E-02 |
| Frem2 | 2.879 | 5.50E-04 |
| Xirp1 | 2.813 | 2.77E-02 |
| Mybpc1 | 2.655 | 1.09E-04 |
| Myl1 | 2.570 | 5.75E-02 |

**B**

**Top Downregulated Genes in 12 Hour Ear Fibroblasts**

| Gene | Expression Change (β) | q-value |
|---|---|---|
| B3galt1 | -2.149 | 8.32E-04 |
| Apc | -1.717 | 9.90E-02 |

**C**

**Top Upregulated Genes in 12 Hour Lung Fibroblasts**

| Gene | Expression Change (β) | q-value |
|---|---|---|
| Clec12a | 2.757 | 2.66E-02 |
| Cd72 | 2.656 | 4.91E-02 |
| Ifi27l2a | 2.397 | 3.05E-04 |
| Ccl12 | 2.255 | 3.05E-04 |
| C1qb | 2.160 | 3.05E-04 |
| Aif1 | 2.103 | 7.71E-02 |
| Rgs18 | 2.062 | 1.67E-02 |
| Nxpe5 | 1.728 | 2.87E-02 |
| Il1rl1 | 1.375 | 3.78E-05 |
| Trpc6 | 1.326 | 7.56E-02 |

**D**

**Top Downregulated Genes in 12 Hour Lung Fibroblasts**

| Gene | Expression Change (β) | q-value |
|---|---|---|
| Nup107 | -4.879 | 4.09E-05 |
| Igsf10 | -2.691 | 1.11E-07 |
| Dnah2 | -2.472 | 5.82E-02 |
| Ctnna2 | -2.369 | 7.71E-02 |
| Tead1 | -1.541 | 1.09E-09 |
| Enpep | -1.508 | 1.75E-02 |
| Mxd3 | -1.487 | 8.14E-06 |
| Fbln1 | -1.446 | 1.09E-09 |
| Mdk | -1.173 | 8.86E-02 |
| Heph | -1.137 | 4.29E-04 |

**E**

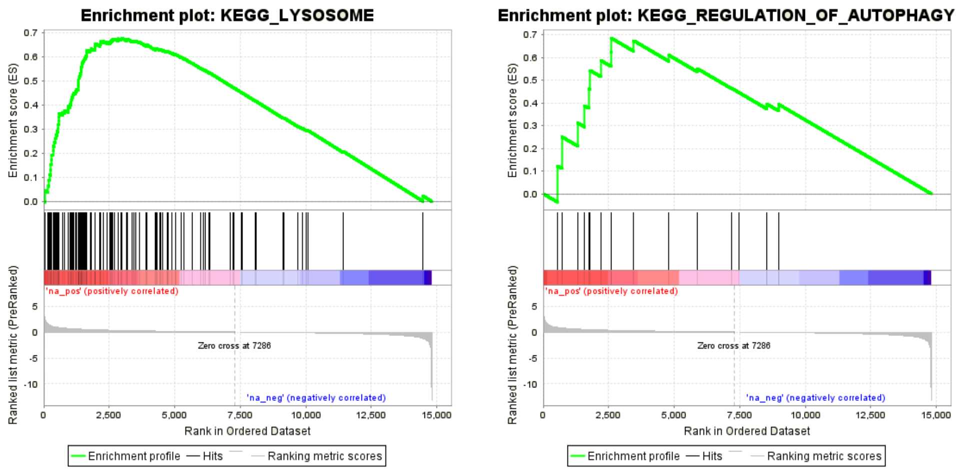

**Figure S8**

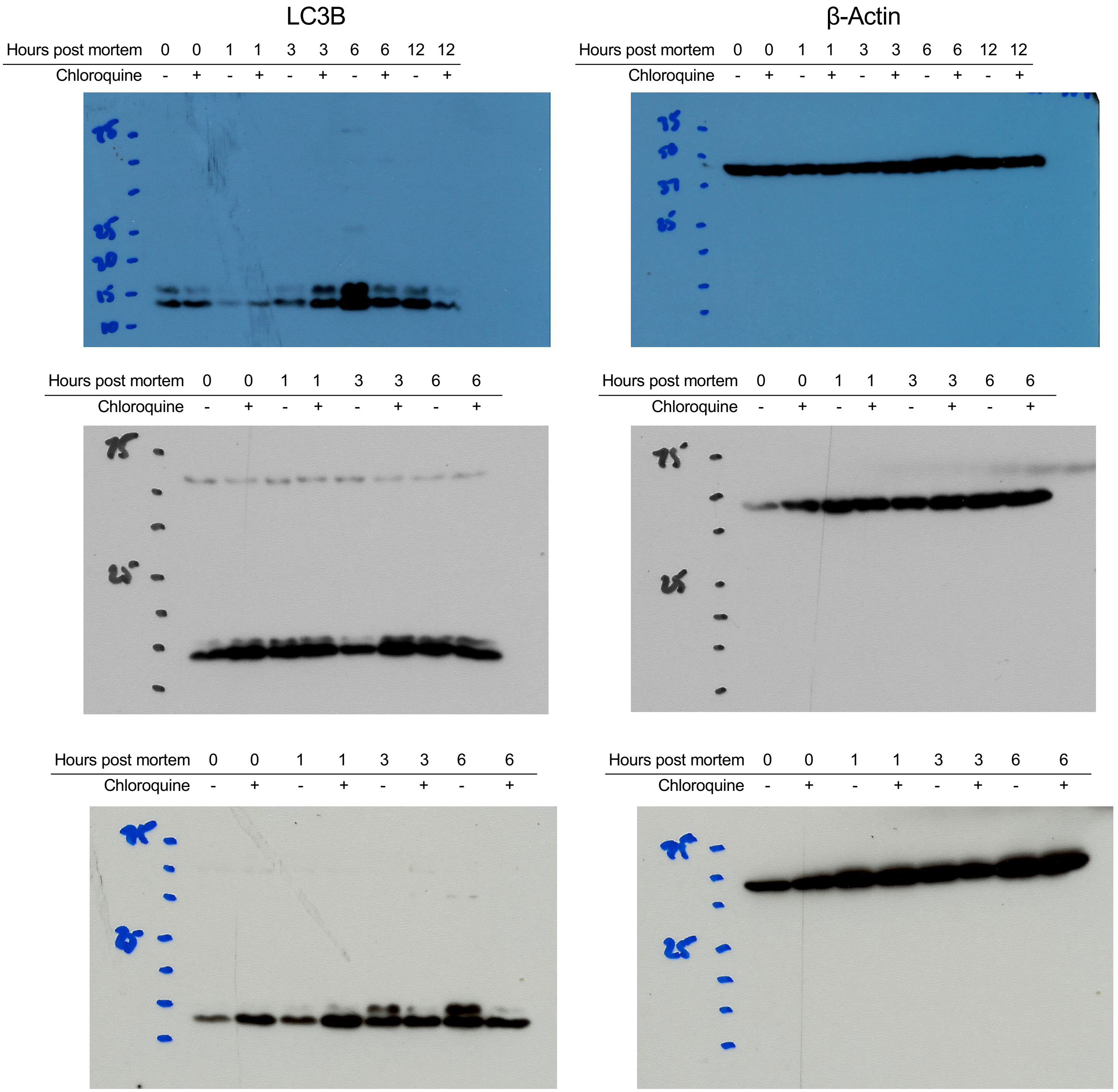
LC3B
β-Actin
Hours post mortem 0 0 1 1 3 3 6 6 12 12
Chloroquine - + - + - + - + - +
Hours post mortem 0 0 1 1 3 3 6 6 12 12
Chloroquine - + - + - + - + - +
Hours post mortem 0 0 1 1 3 3 6 6
Chloroquine - + - + - + - +
Hours post mortem 0 0 1 1 3 3 6 6
Chloroquine - + - + - + - +
Hours post mortem 0 0 1 1 3 3 6 6
Chloroquine - + - + - + - +
Hours post mortem 0 0 1 1 3 3 6 6
Chloroquine - + - + - + - +

**Figure S9**

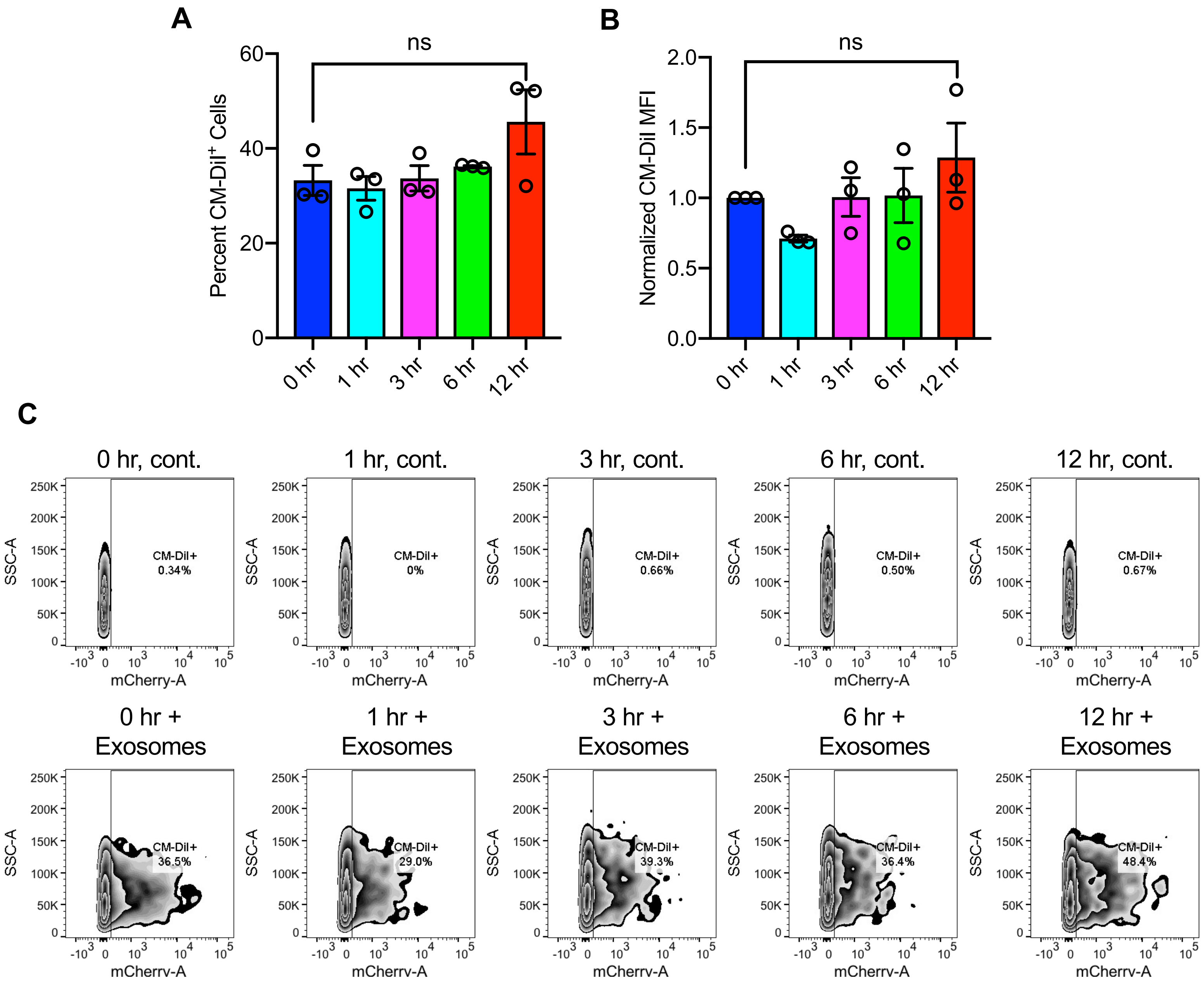
A
ns
Percent CM-DiI+ Cells
60
40
20
0
0 hr
1 hr
3 hr
6 hr
12 hr
B
ns
Normalized CM-DiI MFI
2.0
1.5
1.0
0.5
0.0
0 hr
1 hr
3 hr
6 hr
12 hr
C
0 hr, cont.
CM-DiI+ 0.34%
1 hr, cont.
CM-DiI+ 0%
3 hr, cont.
CM-DiI+ 0.66%
6 hr, cont.
CM-DiI+ 0.50%
12 hr, cont.
CM-DiI+ 0.67%
SSC-A
250K
200K
150K
100K
50K
0
-10^3 0 10^3 10^4 10^5
mCherry-A
0 hr + Exosomes
CM-DiI+ 36.5%
1 hr + Exosomes
CM-DiI+ 29.0%
3 hr + Exosomes
CM-DiI+ 39.3%
6 hr + Exosomes
CM-DiI+ 36.4%
12 hr + Exosomes
CM-DiI+ 48.4%
mCherrv-A
